# Emergent Oscillation Induced by Nutrient-Modulating Growth Feedback

**DOI:** 10.1101/2021.02.09.430447

**Authors:** Juan Melendez-Alvarez, Changhan He, Rong Zhang, Yang Kuang, Xiao-Jun Tian

**Affiliations:** School of Biological and Health Systems Engineering, Arizona State University, Tempe, AZ 85281, USA; School of Mathematical and Statistical Sciences, Arizona State University, Tempe, AZ 85281, USA

## Abstract

Growth feedback, the inherent coupling between the synthetic gene circuit and the host cell growth, could significantly change the circuit behaviors. Previously, a diverse array of emerged behaviors, such as growth bistability, enhanced ultrasensitivity, and topology-dependent memory loss, were reported to be induced by growth feedback. However, the influence of the growth feedback on the circuit functions remains underexplored. Here, we reported an unexpected oscillatory behavior of a self-activation gene circuit induced by nutrient-modulating growth feedback. Specifically, after dilution of the activated self-activation switch into the fresh medium with moderate nutrient, its gene expression first decreases as the cell grows and then shows a significant overshoot before it reaches the steady states, leading to oscillation dynamics. Fitting the data with a coarse-grained model suggests a nonmonotonic growth-rate regulation on gene production rate. The underlying mechanism of the oscillation was demonstrated by a molecular mathematical model, which includes the ribosome allocation towards gene production, cell growth, and cell maintenance. Interestingly, the model predicted a counterintuitive dependence of oscillation amplitude on the nutrition level, where the highest peak was found in the medium with a moderate nutrient but was not observed in rich nutrient. We experimentally verified this prediction by tuning the nutrient level in the culture medium. We did not observe significant oscillatory behavior for toggle switch, suggesting that the emergence of oscillatory behavior depends on circuit network topology. Our results demonstrated a new nonlinear emergent behavior mediated by growth feedback, which depends on the ribosome allocation between gene circuit and cell growth.

## Introduction

Rational design and forward engineering of synthetic gene circuits based on general principles have demonstrated its power in successfully constructing, testing, and debugging circuits for many remarkable applications in the past [1, 2, 3, 4]. However, the complex interplay between the gene circuit and the host cell physiology at various scales often makes the circuit functions fragile to the environmental fluctuations [5, 6, 7, 8, 9, 10]. The transcription and translation of heterologous gene circuits steal a considerable amount of the host cell’s resources that are originally allocated for host activities, thus inevitably leading a significant metabolic burden to the host cell [11, 12, 13, 14]. The altered physiology to host cells, in turn, affects the circuit intended function [15, 16, 17, 18, 19]. Furthermore, the host environmental fluctuations make the circuit-host interplay more complicated and make circuit behavior hard to predict. Thus, it is essential to understand the general principle of how these circuit-host interactions affect the functionality and consider these substantial effects into the design of gene circuits.

One of the most important circuit-host interaction is the growth feedback, in which circuit gene expression slowdowns the host cell growth while the host cell growth also affects the circuit gene expression [20, 21, 22, 23, 18]. Previous studies have reported various emergent circuit properties that result from growth feedback. For example, growth feedback could increase the ultrasensitivity of the gene circuit and thus make a non-cooperative positive autoregulatory system bistable [23]. On the other hand, the growth feedback could also drastically alter circuit functions. Recently, we found a topology-dependent interference of synthetic gene circuit function by growth feedback [18]. We tested two synthetic memory circuits, self-activation switch and toggle switch, and found that both circuits interact with host cells through growth feedback but behave differently. While the self-activation switch quickly lost its memory due to the fast dilution of circuit products mediated by host cell growth, the toggle switch is more robust to the growth feedback and retains its memory after the fast growth phase. It is also reported that an increase of nutrients may lead to a bistable switch circuit transition from bistability to monostability [24]. However, the influence of the circuit dynamics by the growth feedback remains underexplored.

Here we studied how the growth feedback and its effects on gene circuits are modulated by nutrient level. Followed by our previous work [18], we diluted the cells with activated self-activation switch into the fresh medium with reduced nutrition level with a high dose of inducer L-ara. It is surprising to us that the circuit expression level showed very unexpected oscillatory dynamics in a broad range of nutrition levels. Specifically, after dilution into the fresh medium with moderate nutrient, the circuit gene expression first decreases as the cell grows and then shows a significant overshoot before it reaches the steady states. To understand the mechanism of this unexpected phenomenon at the coarse-grained level, we revised our previous model by considering the regulation of the circuit production rate by growth rate. Fitting this model to the experimental data suggests that the production rate could nonmonotically depend on the growth rate. To further understand the mechanism at the molecular level, we built another model by including the ribosome allocation towards gene production, cell growth, and cell maintenance. Interestingly, our model predicted that the oscillation amplitude shows a nonmonotonic relation with respect to the nutrient level, where the highest peak was found in the medium with a moderate nutrient but not a rich nutrient. We experimentally verified this prediction by tuning the nutrient level with various LB fractions in the culture medium. In addition, the emergence of oscillatory behavior depends on circuit network topology as we did not observe significant oscillatory behavior for the toggle switch.

## Results

### Unexpected oscillatory dynamics induced by growth feedback with varied nutrition level

In our previous work, we found that the gene circuits interact with the host cell growth (Fig. 1A), which led to the loss of memory for the self-activation circuit after diluting the activated cells into the fresh 100% Lysogeny broth (LB) rich medium [18]. The memory can be maintained well with the M9 minimal culture medium. Here in this work, we studied how the nutrient level modulates the growth feedback and circuit dynamics (Fig. 1A).

**Figure 1:**
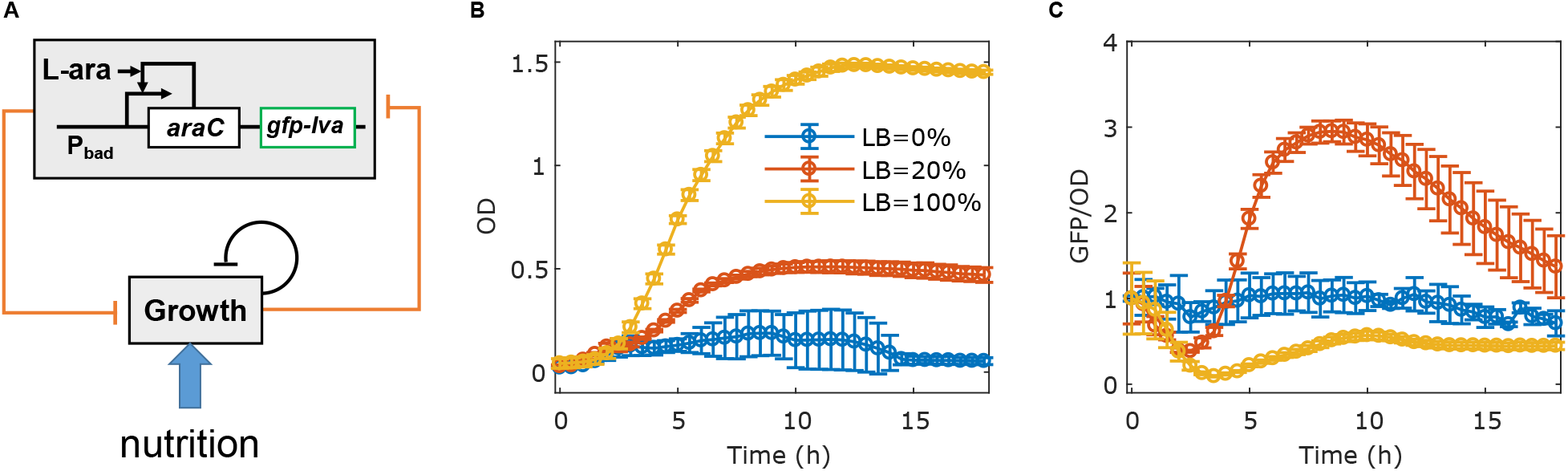
Unexpected oscillatory dynamics induced by growth feedback with varied nutrition level in the SA circuit. (A) Diagram of interactions between the self-activation (SA) gene circuit and the host cell growth, which is modulated by the nutrient. In the SA circuit, transcriptional factor AraC forms a dimer and binds to promoter *P_BAD_* in the presence of stimulus L-(+)-arabinose (L-ara), and thus activates the expression of itself. Unstable GFP variant (GFP-lva) is used as the reporter. (B-C) Dynamics of growth (Optical Density, OD) (B) and GFP/OD (C) after 1:100 dilution of cells with activated SA circuit into fresh medium with a high dose of L-ara and three nutrient levels with 0%, 20% and 100% LB in the culture medium. The data with 100% LB and 0% LB are from previous work [18]. Data indicate mean±s.d. (n=4 biological replicates).

We first varied nutrition levels by mixing 20% of LB media and 80% of M9 media where there is no cell growth. We measured the temporal dynamics of the cell growth and circuit gene expression after diluting the activated cell into a fresh medium with 20% LB and a high/low dose of circuit inducer L-ara. We observed that the cell growth followed the logistic model in both moderate and rich nutrient mediums, with a smaller population carrying capacity in the 20% LB medium (Fig. S1A, Fig. 1B). Consistent with our previous work by diluting cells into the fresh medium with 100% LB and a low dose of L-ara, the circuit memory is also lost due to the high division rate in the medium with 20% LB (Fig. S1B). This result confirmed that the SA circuit is susceptible to growth feedback, even with a moderate nutrient in the growth medium. In the medium with high L-ara, the average GFP level (GFP/OD) decreases through the log-phase until it reaches a minimum, consistent with 100% LB medium (Fig. 1C). However, the circuit dynamics changed dramatically with moderate nutrient after the valley. The GFP level exhibits significant overshoot behavior before the system settles down to the steady state, instead of reaching the steady state directly as in the 100% LB medium. This unexpected oscillation phenomenon suggests that the nutrient could significantly alter the growth feedback and the synthetic gene circuit dynamics.

### Theoretical analysis suggests a nonmonotonic growth-rate regulation on gene production rate

To understand the underlying mechanism of the emerged oscillatory behavior induced by growth feedback with a varied nutrition level, we first tested the simplified coarse-grained model from our previous work [18]. We found that the model is not able to fit the data well (Fig. S2). This is consistent with our theoretical proof that there is no overshoot with this simplified model (See supplementary note, Proposition 1). This simplified model included the growth-mediated dilution but did not consider potential regulations of gene production by growth rate. This model was able to explain the memory loss of the SA circuit after diluting into a fresh medium with rich nutrients where the growth-mediated dilution plays the dominant role, as demonstrated previously [18].

Here the overshoot behavior from the varied nutrient level strongly suggests that host cell growth could increase the production rate of synthetic gene circuits. Thus, we revised the model by adding a growth rate-dependent function *F* (*GR*) on the circuit gene production rate (Fig. 2A, see the Methods for details). We used several different functions to test the monotonicity of *F* (*GR*). We found that both a monotonous function and a linear function of *F* (*GR*) are not able to fit the data well (Fig. 2B-C, red/yellow lines). We further proved that when *F* (*GR*) is a monotonously decreasing function or positive linear function, there is no overshoot (See supplementary note, Proposition 2 and 3).

**Figure 2:**
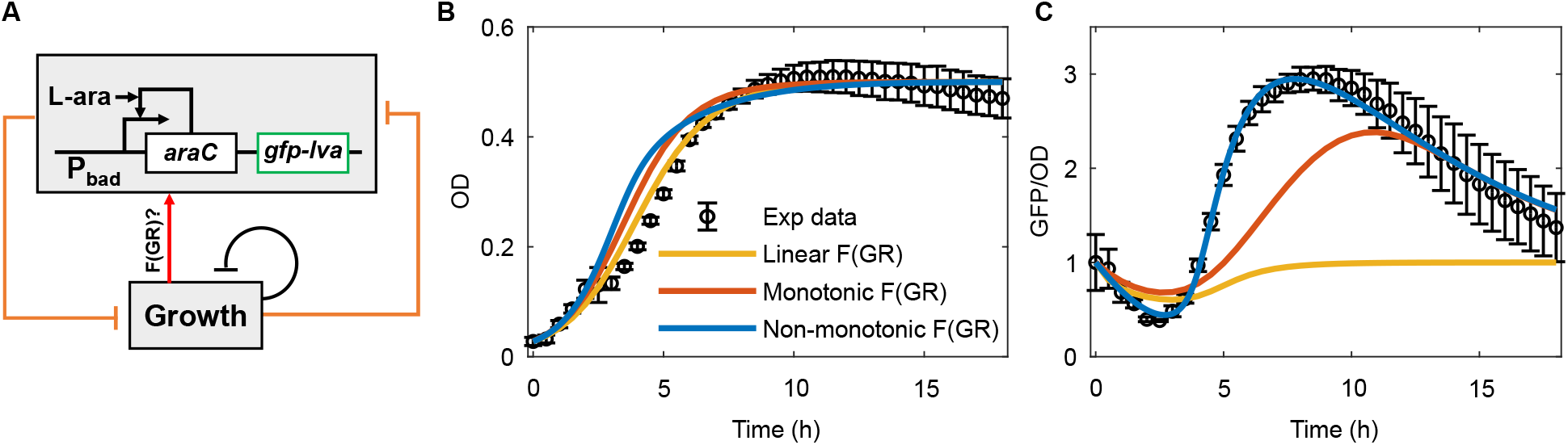
Theoretical analysis suggests a nonmonotonic growth-rate regulation on gene production rate. (A) The diagram of the model by considering the regulation of the production rate of the SA circuit by growth rate. (B-C) Fitting of the model to the dynamics of the host cell growth (B) and the circuit gene expression (C). Linear (yellow lines), monotonic (orange lines), and nonmonotonic (blue lines) functions were used to test the regulation of the production rate of the gene circuit by growth rate (*F* (*GR*)) and only the nonmonotonic function enable the models to fit the experimental data perfectly.

We then used a nonmonotonic function of *F* (*GR*) and found that it fits the data satisfactorily (Fig. 2B-C, blue lines). The fitting result shows that *F* (*GR*) first decreases a bit and then increased with time quickly to a peak and then decreases to the basal level (Fig. S2A). The analysis suggests that *F* (*GR*) first increase and then decrease with growth rate (Fig. S2B). In the early fast growth stage, the reduced production rate and immediate dilution because of fast growth lead to a sharp GFP decrease. As the cell growth slowing down, an increase of the production rate mediated by growth overcome the reduced dilution, leading to the increases and overshoot of GFP. Finally, in the stationary phase, GFP relaxes to the steady state where growth ceases. Taken together, our theoretical analysis suggests that there is a nonmonotonic growth-rate regulation on gene production rate, which accounts for the emergence of the oscillation behavior.

### Molecular mechanism for the growth rate regulation on gene production rate

The regulation of the circuit gene production rate by growth rate is largely dependent on the fact that the ribosome level in the host cell increase with growth rate [20, 25]. To further understand the oscillatory behavior at the molecular level, we also developed a mathematical model by integrating the ribosome dynamics and its allocation to the gene production and gene expression for the host cell growth (Fig. 3A) [see method for detail). Basically, the ribosome level increases with the growth rate via an autocatalysis way. The allocation of ribosome to host cell growth and synthetic gene expression depends on the compromised growth rate and resource demands.

**Figure 3:**
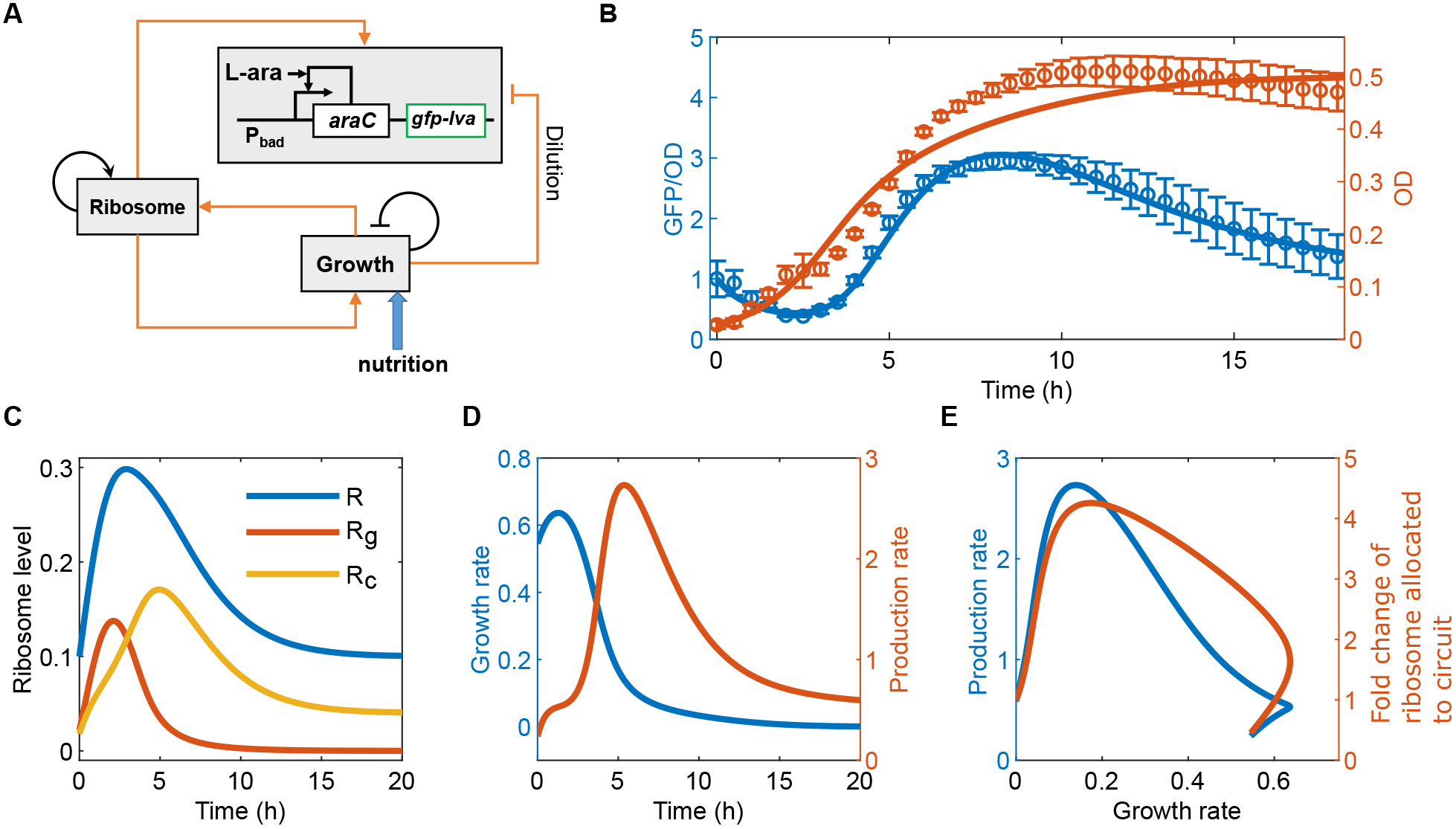
Molecular mechanism for the growth rate regulation on gene production rate. (A) The diagram of the model by considering the ribosome regulation and allocation. (B) Fitting of the model to the dynamics of the host cell growth and circuit gene expression. Solid lines are model simulation, and the dotted line with error bars are experimental data. (C) The temporal dynamics of the ribosome production and allocation to the cell growth and the SA circuit. (D) The temporal dynamics of the growth rate and production rate of the synthetic gene circuit. (E) The relation between the production rate and fold change of ribosome allocated to the synthetic circuit and growth rate.

Upon data fitting, this model successfully captured the oscillatory dynamics of the circuit gene expression (Fig. 3B). The ribosome is increased quickly after the dilution but is mainly allocated to cell growth at the early stage (Fig. 3C, blue and red lines). The allocated ribosome to the circuit is limited at this stage (Fig. 3C, yellow line), and thus the production of the gene circuit is kept at a low level (Fig. 3D, blue line). The limited ribosome to the circuit and the dilution from the fast growth make the GFP level decreasing quickly. When the cell growth becomes slower, the ribosome level has accumulated and continues to increase due to the autocatalysis (Fig. 3C, black line), but the ribosome demand from the cell growth starts to decrease (Fig. 3C, red line), thus leading to the spare ribosome allocated to the gene circuit (Fig. 3C, blue line), a fast increase of the production rate of the circuit gene expression (Fig. 3C, yellow line), and the overshoot of GFP level to a very high level. When the ribosome decreases to the low level at the stationary phase, the GFP level also reaches the steady-state. The relationship between the circuit gene expression rate and the growth rate follows a nonmonotonic curve (Fig. 3E), consistent with the suggestion in the last section. Thus, the model not only provides a molecular mechanism for the emerged oscillation induced by growth feedback but also for nonmonotonic growth-rate regulation on gene production rate.

### The amplitudes of the oscillatory dynamics are controlled by nutrient level

Fig. 1 suggested that the oscillation is emerged in some range of nutrient level but vanished after the one nutrient threshold. To systematically study how the oscillatory behavior depends on the nutrient level, we analyzed the dynamics of the circuit gene expression by varying one parameter, carrying capacity (*N_max_*), in the model. *N_max_* is a good indicator of the nutrient level, given that the cell density reaches differently carrying capacity according to the nutrient level. It is noted that even *N_max_* is not proportional to nutrient level, the positive dependence of *N_max_* on the nutrient level could still give us a qualitative prediction about how the behavior is emerged and vanished with the nutrient levels.

Figure 4A shows the model prediction about the dynamics of the circuit gene by varying *N_max_* systematically. It is clear that the circuit gene expression always decreases immediately after dilution into the fresh medium to a valley (Fig. 4A), and the depth of the valley increases with *N_max_* (Fig. 4B). This is reasonable given that the larger *N_max_* gives a longer period of fast growth and thus the more dilution of the circuit gene expression, consistent with our previous work. After reaching the valley, the system starts to bounce back to the steady state with an overshoot and shows a distinct oscillatory behavior for *N_max_* < 1.15, or directly relaxes to the steady state without overshoot and shows an adaptation behavior if *N_max_* > 1.15. That is, the oscillation can be emerged with moderate nutrient levels but vanishes with rich nutrient. Interestingly, the oscillation peak first increases to a maximum at *N_max_* < 0.379, and then decreases with *N_max_* until converging with the steady state (Fig. 4B). The transition from the oscillation to the adaptation largely depends on ribosome allocation and the gene activity at the valley points. Large *N_max_* creates a long period of fast growth, which dilutes the circuit gene expression significantly and makes more ribosome allocated to cell growth (Fig. S3A-C). Furthermore, for large *N_max_* at the valley point, the low level of AraC leads to a very weak production of gene circuit due to its self-activation topology (Fig. S3D). Thus the circuit is not able to take advantage of the spared ribosome for overshooting before the system reach to steady state (Fig. S3C).

**Figure 4:**
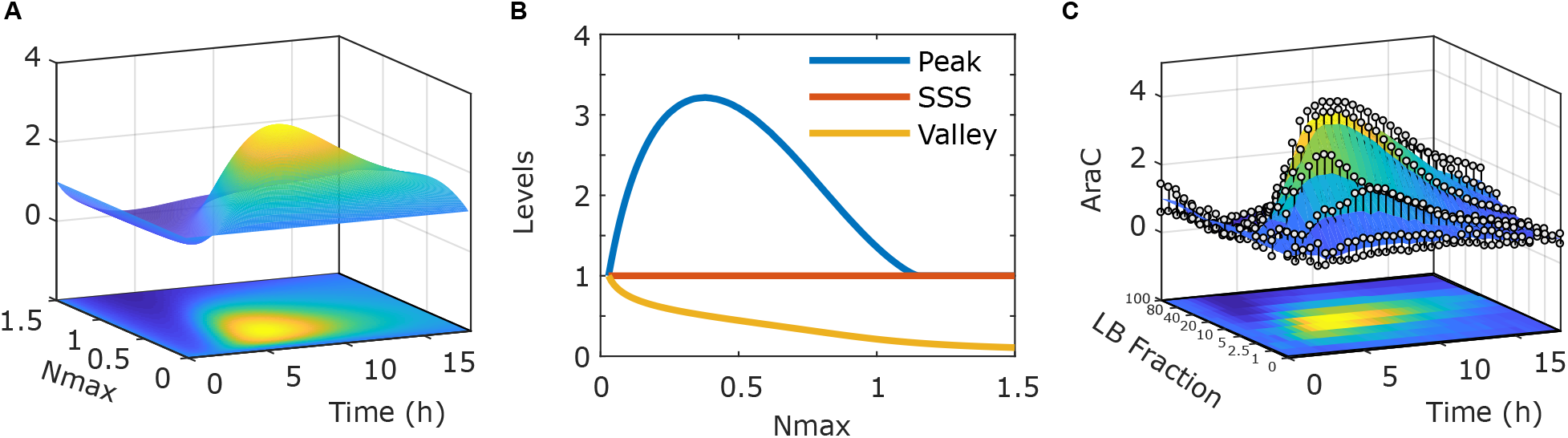
The amplitudes of the oscillatory dynamics in the SA circuit are controlled by nutrient level. (A-B) Model prediction about the dynamics of gene expression level (A) and the amplitudes of the oscillations (B) with various carrying capacity (*N_max_*) in the SA circuit. (C) Dynamics of GFP/OD after 1:100 dilution of cells with activated SA switch into the fresh medium with various LB fractions. Data indicate mean±s.d. (n=4 biological replicates).

To verify this prediction experimentally, we tuned the nutrient level by varying the LB fraction in the culture medium from 0 to 100%. After dilution of the cells into the fresh medium, the host cell growth followed the logistic model to reach different carrying capacity according to the fraction of LB in the culture medium (Fig. S4A). The dependence of carrying capacity (*N_max_*) on the LB fraction shown in Fig. S4B confirmed that the *N_max_* is a good indicator of the nutrient level in the theoretical analysis in Fig. 4AB. The GFP dynamics with various LB fractions is shown in Fig. 4C. A general decrease of GFP immediately after the dilution was observed for all the nutrient levels, with the depth increasing with nutrient level, consistent with the model prediction. Most importantly, oscillatory behavior was found in a broad nutrient range with 1%∼ 40% LB, where the maximum peak was found in the medium with 10% LB. Overshoot was not found in the medium with 80% and 100% LB, where the GFP level was relaxed to the steady state after the valley point. Thus, we experimentally validated our modeling prediction.

### The emergence of oscillatory behavior depends on circuit network topology

To further investigate whether the emergent oscillation also exists in the toggle switch, which is is another well-characterized bistable circuit with two genes (LacI and TetR) repressing the expression of each other (Fig. 5A). In our previous work, we compared the SA switch and the toggle switch circuits in response to growth feedback. We found that the SA circuit is very sensitive to the growth-mediated dilution and lost its memory very easily, while the toggle switch was more refractory to the growth feedback and retrieved its memory after the fast-growth phase[18]. Here we tested whether these two gene circuits behave differently in response to the nutrient-modulating growth feedback.

**Figure 5:**
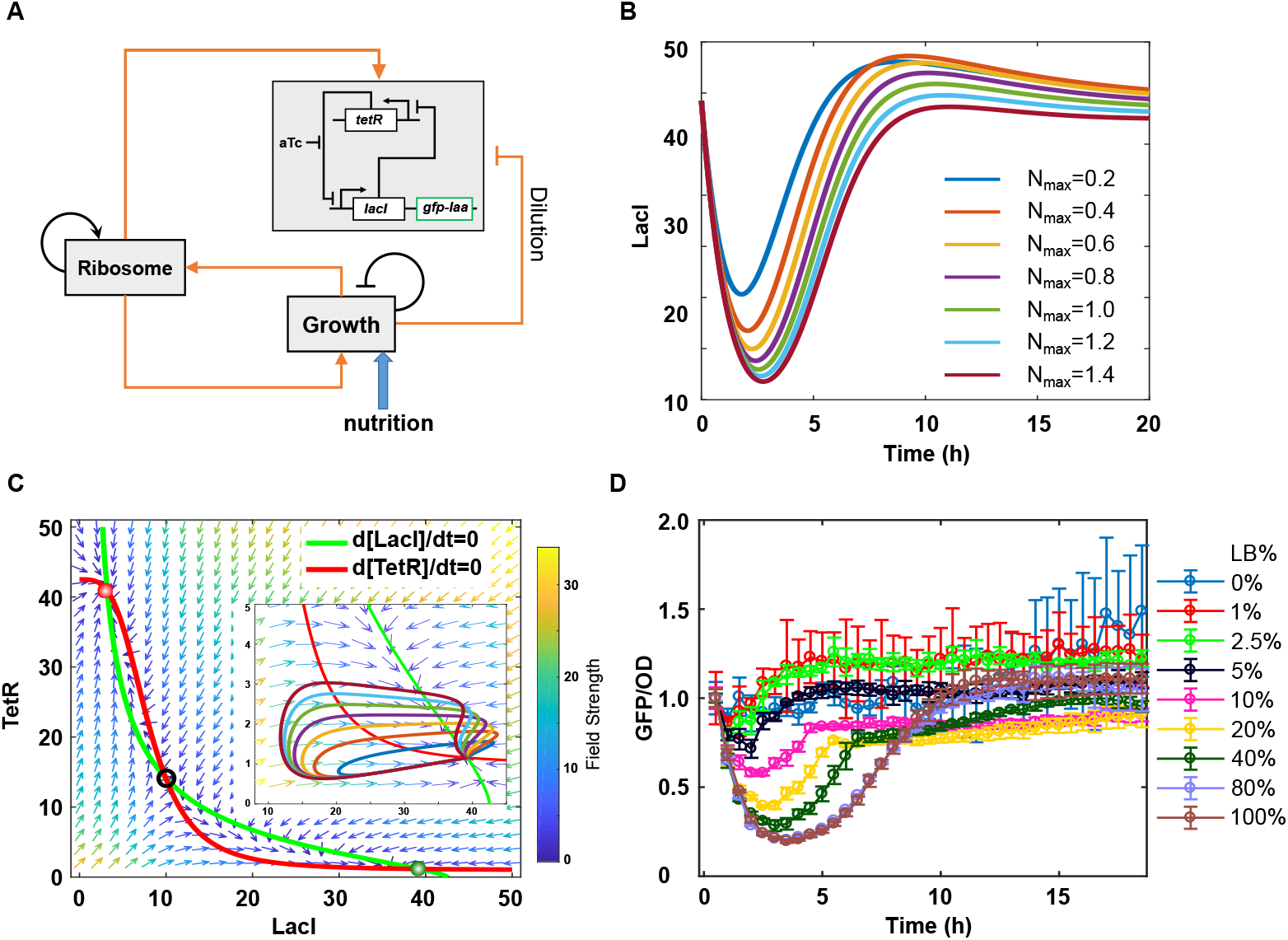
The emergence of oscillatory behavior depends on circuit network topology. (A) The diagram of the model for toggle switch circuit by considering the ribosome regulation and allocation. (B-C) Model prediction about the temporal dynamics of circuit gene expression level (B) and trajectories on the phase plane diagram (C) with various carrying capacity (*N_max_*). The nullclines of LacI and TetR are shown in green and red, respectively. The vector field of the system is represented by small arrows and the field strength is indicated by the color of the arrows. (D) Dynamics of GFP/OD after 1:100 dilution of cells with activated toggle switch into fresh medium with various LB fractions. Data indicate mean±s.d. (n=3 biological replicates).

We applied the same modeling framework to the toggle switch (Fig. 5A, and method for details). We did a theoretical analysis on the dynamics of the toggle switch after dilution of the host cells into the fresh medium by varying the carrying capacity *N_max_*. We found that the LacI level dropped quickly to different levels according to the *N_max_* values (Fig. 5B). It is noted that the LacI level at the valley point decreases with *N_max_* (Fig. 5B), similar to the SA circuit (Fig. 4A). This is reasonable because larger *N_max_* ensures a longer duration of fast-growth phase and thus longer dilution-dominated phase. Interestingly, we did not observe a significant overshoot for the toggle switch (Fig. 5B), which is different from the SA circuit. To illustrate the underlying mechanism, we analyzed the simulated trajectories in the direction field. As shown in Fig. 5C, the system was initially in the LacI-high state (Green circle) and then moved toward LacI-low/TetR-low corner due to the dilution of both LacI and TetR in the fast-growth phase. Then, the system recovered back to its original state following the direction field without significant overshoot for all nutrient levels.

Following the same experimental protocol to verify the prediction, we diluted the cells with activated toggle switch into to fresh medium with different LB fractions. We found that after dilution the GFP level decreased to a minimum that depends on the LB fraction (Fig. 5D), consistent with the model prediction. Furthermore, no significant overshoot was observed for all samples with various LB fractions (Fig. 5D), which is also consistent with the model prediction. Thus, the emergence of oscillatory behavior depends on circuit network topology.

## Discussion

Here, we simply tuned the growth feedback by modulating the nutrient level with mixed LB and the M9 minimal culture media. In the future, it would be interesting to use a chemically-defined medium with known chemical composition to fine-tune the growth feedback and study its effects on gene circuits systematically. Recently, it is found that protein production strategies differ in response to starvation for carbon, nitrogen, or phosphate [26, 27]. However, it is unclear how these different strategies regulations under these nutrient limitations affect the growth feedback and the synthetic gene circuits. In addition, a fluctuating growth environment in different host cells affects the host cell metabolism and the circuit functions [28, 29]. Systematical measurement of the perturbed transcriptional/translational rates, growth rate, and the ribosome profiles will significantly enhance our understanding of the modulation of growth feedback by nutrient levels and limits.

Understanding the growth feedback at the molecular level will greatly help us to formulate control strategies for engineering gene circuits robust to the host cell physiological environment. Here we considered the ribosome allocation in our mathematical model and found that the interaction between ribosome availability and growth condition due to different nutrient levels can produce unexpected outcomes in synthetic gene circuit dynamics. The competition of cellular resources between the endogenous genes and heterologous synthetic genes exist across multiple levels, including RNA polymerase (RNAP) at the transcriptional level and ClpXP at the protein turnover level, which are not considered here. Furthermore, it is unclear whether there is a priority for resource allocation.

It is reported that guanosine tetraphosphate (ppGpp) plays a critical role in regulating bacterial growth, and ribosome regulation and allocation [30, 31, 32, 33, 34, 35]. Sophisticated regulations have been found on the regulations of ribosome and ppGpp, such as feedforward loop [36], positive feedback [37]. Interestingly, ppGpp plays a central role in coordinating transcription and translation under nitrogen starvation, but dispensable under carbon starvation [26]. Integrating all of these orchestrators into one modeling framework will provide us a comprehensive understanding of growth feedback.

The effects of growth feedback on the gene circuits are diverse and depend on many factors. The effects could be beneficial and thus can be utilized to enhance the circuit functions, such as the increased ultrasensitivity [23]. The effects could also be detrimental, such as the loss of memory in the self-activation switch [18]. All these focused on the perturbation of the circuit steady-state behaviors by growth feedback. Here, we reported the unexpected alteration of the circuit dynamics by growth feedback. While the gene expression in the SA switch circuit shows emergent oscillation behavior with undershoot and overshoot around the steady state level, the gene expression in the toggle switch circuit only shows undershoot. This is another evidence that the toggle switch is more robust to growth feedback than the SA switch. Characterization of the general principles for the kinetic changes due to fluctuating growth conditions will facilitate our ability to utilize or minimize the effects of growth feedback.

## Methods and Models

### Strains, media, chemicals and plasmids

*E. coli* DH10B (Invitrogen, USA) was used for all the plasmid preparations. Measurement of the self-activated circuit was performed in *E. coli* K-12 MG1655ΔlacIΔaraCBAD, and measurement of the toggle switch was performed with *E. coli* K-12 MG1655ΔlacI as described in [18]. The culture media for the cells were LB broth (Luria-Bertani broth, Sigma-Aldrich) or LB plates supplemented with 25 *μ*g/ml chloramphenicol or 50 g/ml kanamycin depending on the backbone of the circuit. Inducers L-ara (L-(+)-Arabinose and aTc (Anhydrotetracycline hydrochloride, Abcam) were dissolved in ddH2O at concentrations of 25% and 1mg/ml, and stored at −20°*C* in aliquots as stocking solutions. The aTc stocking solutions were replaced every month. When diluted into appropriate working solutions in ddH2O, L-ara solutions were replaced monthly, and aTC solutions were prepared freshly each time and discarded after 24 hours. All the working solutions were kept at 4°*C* and added into culture media with 1000-fold dilution. Details of the self-activated circuit and toggle switch can be found in [18].

### Circuit inductions

The experimental procedure for each biological replicate of the self-activated circuit (SA-circuit) induction was carried out like this. On day one, SA-circuit plasmid was transformed into *E. coli* strain K-12 MG1655ΔlacIΔaraCBAD which were grown on LB plate with 50 *μ*g/ml kanamycin overnight at 37°*C*. On day two in the morning, one colony was picked and inoculated into 400 *μ*l LB medium with 25 g/ml chloramphenicol and was grown to OD 1.0 (measured in 200 *μ*l volume in 96-well plate by plate reader for absorbance at 600 nm) in a 5 ml culture tube in the shaker. The cells were then diluted 1000 folds into 2 ml fresh culture medium supplemented with desired concentration of L-ara, and grew in a 15 ml culture tube with 250 revolutions per minute at 37°*C* for 16 hours. On day three, cells inducted in last step were 100-fold diluted into each culture medium of the nutrient gradient supplemented with desired concentration of L-ara and antibiotics. Then, three technical replicates of 200 *μ*l culture mix for each component of the nutrient gradient were load onto a 96-well plate, which was immediately placed onto the plate-reader to start the measurement.

The experimental procedure for each biological replicate of the toggle switch induction was carried out as above, except for the strain being K-12 MG1655ΔlacI, antibiotics being 50*μ*g/ml kanamycin and the inducer being aTC.

### Dynamic analysis performed by Plate Reader

Synergy H1 Hybrid Reader from BioTek was used to perform the average fluorescence analysis. 200 *μ*l of culture was loaded into each well of the 96-well plate. M9 broth without cells was used as blank. LB broth without cells was used to set the low-scale of the fluorescence. The plate was incubated at 37°*C* with orbital shaking at the frequency of 807 cpm (cycles per minute). The duration of the measurement was 18 hours and the interval between each measurement was 30 minutes. Optical density (OD) of the culture was measured by absorbance at 600 nm; GFP was detected by excitation/emission at 485/515 nm.

### Mathematical model for the SA circuit by considering the growth-rate regulation on gene production rate

Here, we followed our previous work [18] to model the dynamics of the circuit gene expression and cell density. For this work, the dependence of the production rate of the gene circuit on the growth rate of the host cell is considered, following the suggestion about the realistic description of growth effects [38]. The revised model follows is given by:

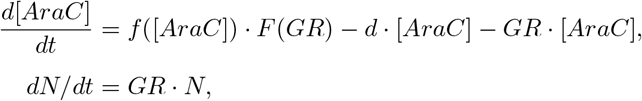

where *f* = *k*_0_ + *k*_1_ · (*S_a_* · [*AraC*]^2^)*/*(*S_a_* · [*AraC*]^2^ + 1), *k*_0_ and *k*_1_ are is the basal production rate and the maximum L-ara-induced production rate of AraC respectively, *S_a_* describes how the production rate is regulated by the inducer L-ara, *d* and *GR* are the degradation rate and dilution rate respectively. [*AraC*] is the concentration of AraC, which is co-expressed with GFP and thus used interchangeably. *N* is the cell density, the growth rate *GR* = *k_g_* · (1 − *N/N_max_*)*/*([*AraC*]*/J* + 1), where *J* is defined as the overload parameter of the gene circuit to the growth rate. *kg* is the maximal growth rate and *N_max_* is the carrying capacity. Here we considered several functions of *F* (*GR*) to fit the experimental data and to find the correct phenomenological dependence of the synthetic gene production rate on growth rate. The function *F* (*GR*) represents the regulation of synthesis rate of AraC by the growth rate. We used *g*(*GR*) = 1 to describe the scenario where the synthetic gene production rate is independent on growth rate, *F* (*GR*) = *a*·*GR*+*b* (linear), and *F*(*GR*) = (*a·GR+b*)*/*(*c·GR+1*) (monotonic) to describe the scenario where the synthetic gene production rate monotonically increases with growth rate, and 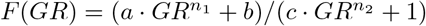 (nonmonotonic, with *n*_1_ < *n*_2_) to describe the scenario where the synthetic gene production rate nonmonotonically changes with growth rate. The best fitted parameters with this model, unless otherwise mentioned, are, *a* = 0.0079, *b* = 1, *k*_0_ = 0.4488, *k*_1_ = 8.9770, *S_a_* = 1, *d* = 4.4885, *k_g_* = 0.9634, *J* = 2.8066; for the linear case. While for the monotonic case, are, *a* = 727.7, *b* = 1, *c* = 277.5 *k*_0_ = 0.0289, *k*_1_ = 0.5779, *S_a_* = 1, *d* = 0.2889, *k_g_* = 0.9065, *J* = 8.1291; and nonmonotonic case are *a* = 7.549, *b* = 1, *c* = 6.7002, *n*_1_ = 0.5, *n*_2_ = 2, *k*_0_ = 0.0514, *k*_1_ = 1.0288, *S_a_* = 1, *d* = 0.5144, *k_g_* = 1.2540, *J* = 2.1230.

### Mathematical model for the SA circuit by considering the ribosome allocation to the host cell growth and synthetic gene circuits

In order to understand the underlying molecular mechanism for the oscillatory dynamics, we also developed a model by explicitly considering the ribosome allocation for cell maintenance, growth feedback, and circuit gene. The ribosome is considered as a primary determinant of growth rate [39]. The revised equation for the gene circuit follows

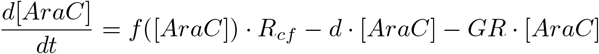

where *R_cf_* is the relative ribosome allocated to the synthetic gene circuit, which depends on the cellular ribosome level and growth rate (see below).

The ribosome level is not a constant but heavily depends on the cell growth rate. It has found that the *E. coli* cell tunes its ribosome level to be linearly correlated with growth rate [40, 20, 41]. Furthermore, at zero growth, the ribosome level in *E. coli* cells is maintained at a basal level [40, 27], which supports the normal physiology of cell with a constant protein production activity in the stationary phase for a very long time [42] and could also support the gene expression in synthetic gene circuits [18]. In addition, the production of the ribosome is also regulated by its autocatalysis [43, 44]. Here we incorporated these regulations into the mathematical model to represent the dynamic regulation of ribosome,

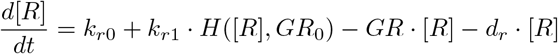

where, we set *H*([*R*], *GR*_0_) = *GR*_0_ · [*R*]^*n*^/([*R*]^*n*^ + *K^n^*) to describe that the ribosome synthesis is linearly controlled by growth rate and also its autocatalysis. Here we used a Hill function to describe the autocatalysis of ribosomes. This is consistent with the experimental observation that ribosomal promoter activity increases with growth rate [41]. By solving this equation at zero growth (*GR*_0_ = 0), we can get the basal level of ribosome as *R_min_* = *k_r_*_0_*/d_r_*.

Here, the ribosome is allocated into three classes for simplicity, including cell maintenance (*R*_0_), Growth (*R_g_*) and circuit gene production (*R_C_*). First, we assume that a constant amount of ribosome *R*_0_ is allocated to cell normal maintenance with a high priority for translating the genes that are essential to the cell survival, including house-keeping genes. Then the rest of ribosome could be allocated to cell growth if the nutrient is sufficient or synthetic gene circuits according to their demands. The production of the exogenous genes can cause a metabolic burden to the host [18]. The growth rate without the metabolic burden is represented by *GR*_0_ = *k_g_* · (1 − *N/N_max_*). The ribosome working on growth is given by *R_g_* = (*GR*_0_*/GR*) · (*R* − *R*_0_). The ribosome expression at zero growth rate (*R_min_*) is able to maintain a steady state level of the gene expression. The amount of ribosome producing AraC at steady state is (*R_min_* − *R*_0_). Therefore, the fraction of ribosome available for gene production in respect to steady state of the circuit is represented by *R_cf_* = (*R* − *R_g_* − *R*_0_)*/*(*R_min_* − *R*_0_). The best fitted parameters with this model, unless otherwise mentioned, are, *k*_0_ = 0.0335, *k*_1_ = 0.6698, *S_a_* = 1.1453, *d* = 0.3349, *k_g_* = 0.9945, *J* = 2.1919, *k_r_*_0_ = 0.0879, *k_r_*_1_ = 1.7581, *K* = 0.7988, *d_r_* = 0.8790, *n* = 1 and *R*_0_ = 0.0599.

### Mathematical model for the toggle switch

Following the model for the toggle switch in our previous work [18, 29] and the above modeling framework for the SA circuit, we developed the following ODE model to describe the dynamics of LacI and TetR concentration ([LacI] and [TetR]) in the toggle switch,

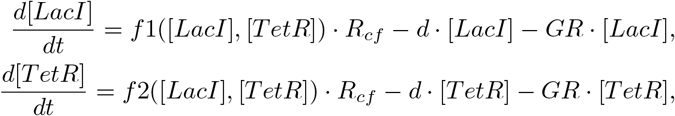

where 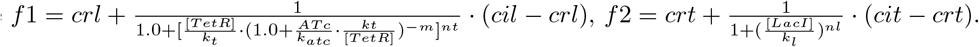 *crl* and *cil* are the production rates of LacI when its promoter is repressed or induced, respectively. *k_t_* is the TetR concentration that makes 50% of the maximum inhibition on LacI, and *nt* describes the ultrasensitivity of this inhibition. *m* is the Hill coefficient to describe the relationship between the active ratio of repressor TetR and the aTc inducer concentration. *crt* and *cit* are the production rates of TetR when its promoter is repressed or induced. *k_l_* is the LacI concentration that makes 50% of the maximum inhibition on TetR, and *nl* describes the ultrasensitivity of this inhibition. *d* is the degradation rate constant, and the *GR* is growth rate, which is represented by *GR* = *k_g_* · (1 − *N/N_max_*)*/*(1 + ([*LacI*] + [*TetR*])*/J*). LacI is co-expressed with GFP, and thus was used interchangeably. *R_cf_* can be calculated following the modeling framework in the above section for the SA circuit. The best fitted parameters with this model, unless otherwise mentioned, are, *crl* = 0.1, *cil* = 4.25, *k_t_* = 6, *k_atc_* = 0.54, *m* = 3, *nt* = 1.5, *crt* = 0.5, *cit* = 4.25, *k_l_* = 8, *nl* = 3.5, *d* = 0.1, *J* = 50, *k_g_* = 1.2, *ATc* = 0, *k_r_*_0_ = 0.0879, *k_r_*_1_ = 1.7581, *K* = 0.7988, *d_r_* = 0.8790, *n* = 1 and *R*_0_ = 0.0599.

### Parameter fitting

For fitting of parameter to the experimental data, we used the Matlab function fminsearch, which minimizes the least square error between the model simulation results and the experimental data. With the purpose of following experimental designs, we set the model initial conditions to the high expressed GFP level steady state.

## Data availability

All data produced or analyzed for this study are included in the published article and its supplementary information files or are available from the corresponding author upon reasonable request.

## Code availability

All the equations and parameters of the mathematical models can be found in the supplementary note.

## Author Contributions

X-J.T. conceived the study. X-J.T., R.Z., and Y.K. designed the study. R.Z. performed experiments. J.M-A., C.H, Y.K., and X-J.T. performed modeling analysis. R.Z., J.M-A., and X-J.T. analyzed the experimental data. J.M-A. and X-J.T. wrote the manuscript. C.H., R.Z., and Y.K. edited the manuscript.

## Acknowledgments

This project was supported by the ASU School of Biological and Health Systems Engineering and NSF grants (EF-1921412 to X-J.T.;DEB-1930728 to Y.K), NIH grant (5R01GM131405 to Y.K.). J.M-A. were also supported by the Arizona State University Dean’s Fellowship.

## Supplementary Materials

### Mathematical proof on the monotonicity of the growth rate-dependent function on the circuit gene production rate

The general form of the system is:

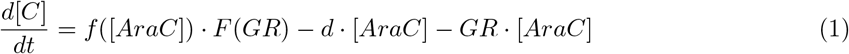

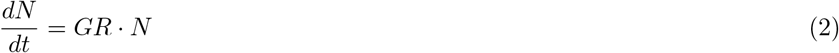

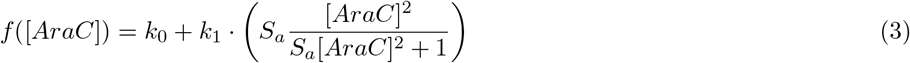

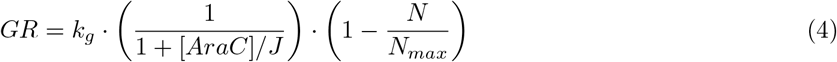

where function *F* can vary.

For simplification, we introduce following notations:

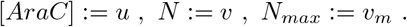

Thus the original model can be written as:

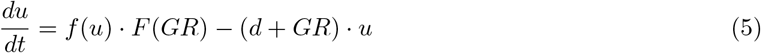

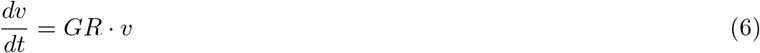

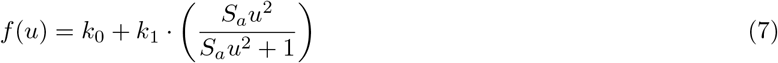

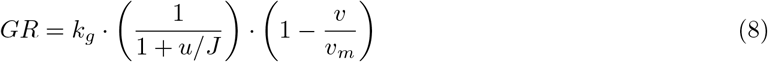

#### Proposition 1

There is no peak of AraC (i.e. no local maximum point of *u* with *t* ∈ (0, *T*) where *T* is the ending time) when *F* (*GR*) = 1. This is corresponding to our previous simplified model, where the regulation of the production rate by the growth rate is not considered [18].

**Proof**: Prove by contradiction. Assuming there is at least one local maximum point *u_p_* = *u*(*t_p_*) with *t_p_* ∈ (0, *T*), which leads to 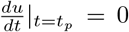. Thus, there exist *t*_1_, *t*_2_ close to *t_p_* such that *t*_1_ < *t_p_* < *t*_2_ and *u*_1_ = *u*_2_ < *u_p_* (Notice that here we denote *u*(*t*_1_):= *u*_1_ and *u*(*t*_2_):= *u*_2_, similar notation for *v*_1_ and *v*_2_) and

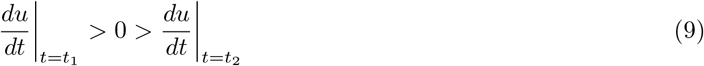

which is equivalent to

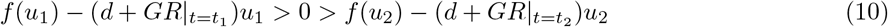

Since *f* (*u*) and *du* are monotonous about *u*, *f* (*u*_1_) = *f* (*u*_2_) and *du*_1_ = *du*_2_ hold, which means the following inequality should also hold:

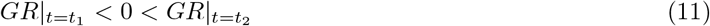

On the other hand, since *GR* is non-negative, *v* is non-decreasing over time *t* and *v*_1_ ≤ *v*_2_ holds, which bring us

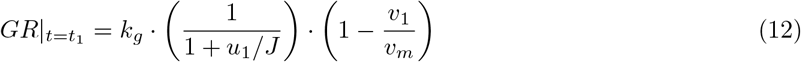

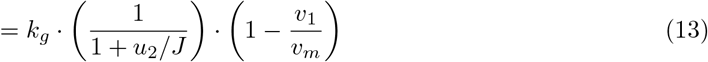

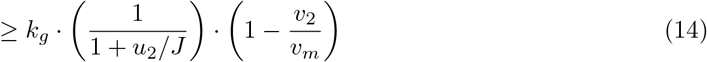

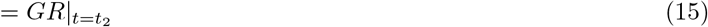

This conflicts with Eq. 11.

#### Proposition 2

There is no peak when *F* (*GR*) is a monotonous decreasing function.

**Proof**: Similarly, we assume there is at least one local maximum point *u_p_* with *t_p_* ∈ (0, *T*),, 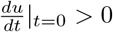 and we prove by contradiction. Thus, there exist *t*_1_, *t*_2_ close to *t_p_* such that *t*_1_ < *t_p_* < *t*_2_ and *u*_1_ = *u*_2_ < *u_p_*. And also we have:

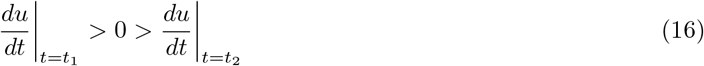

However, since *v*_1_ ≤ *v*_2_ we have

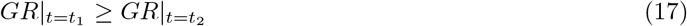

so

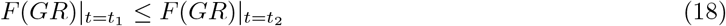

which provides us

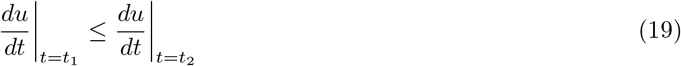

which is conflicting with Eq. 16.

#### Proposition 3

If *F* (*GR*) = *a* · *GR* + 1 where *a* is a positive constant. Then the following two situations will not occur at the same time:

(i). there is a local minimum point *u_v_* with *t_v_* ∈ (0, *T*) and *u_v_* < *u*_0_;
(ii). there is a local maximum point *u_p_* with *t_p_* ∈ (0, *T*) and *u_p_* > *u*_0_; here *u*_0_ = *u*(0) is the initial concentration.

**Proof**: We prove by contradiction. Assume the above two situations occur at the same time, which means there is at least one local minimum point and at least one local maximum point. We assume that among all these extrema, the first one is a local minimum which denoted as *u_v_* with *t_v_* ∈ (0, *T*) and *u_v_* < *u*_0_ (Otherwise we will have similar proof). And we denote the local maximum following *u_v_* as *u_p_* with *t_p_* ∈ (0, *T*) and *u_p_* > *u*_0_. Thus, we have 0 < *t_v_* < *t_p_* < *T* and *u_v_* < *u*_0_ < *u_p_*.

So there exists time *t*_∗_ where *t*_∗_ ∈ (*t_v_, t_p_*) such that *u*(*t*_∗_):= *u*_∗_ = *u*_0_ and

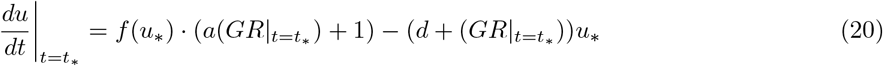

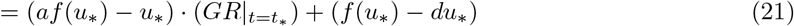

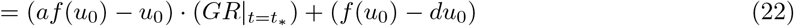

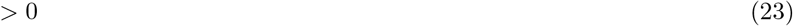

Since *u*_0_ is a steady state of the system without growth-mediated feedback (i.e. *GR* = 0), we have *f* (*u*_0_) − *du*_0_ = 0 which leads to:

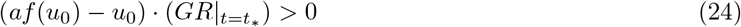

Meanwhile, since *u_v_* is the first extremum, we have

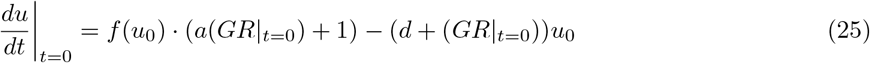

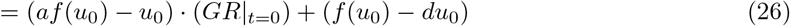

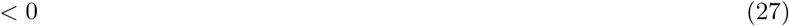

However, from Eq. 24 we know that 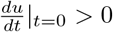, which is a contradiction.

Based on the above propositions we know that, a model can not capture the experimental observations if function *F* (*GR*) is in one of the following forms:

(1). *F* (*GR*) = 1;
(2). *F* (*GR*) is a monotone decreasing function;
(3). *F* (*GR*) is a linear function.

---

**Figure S1:**
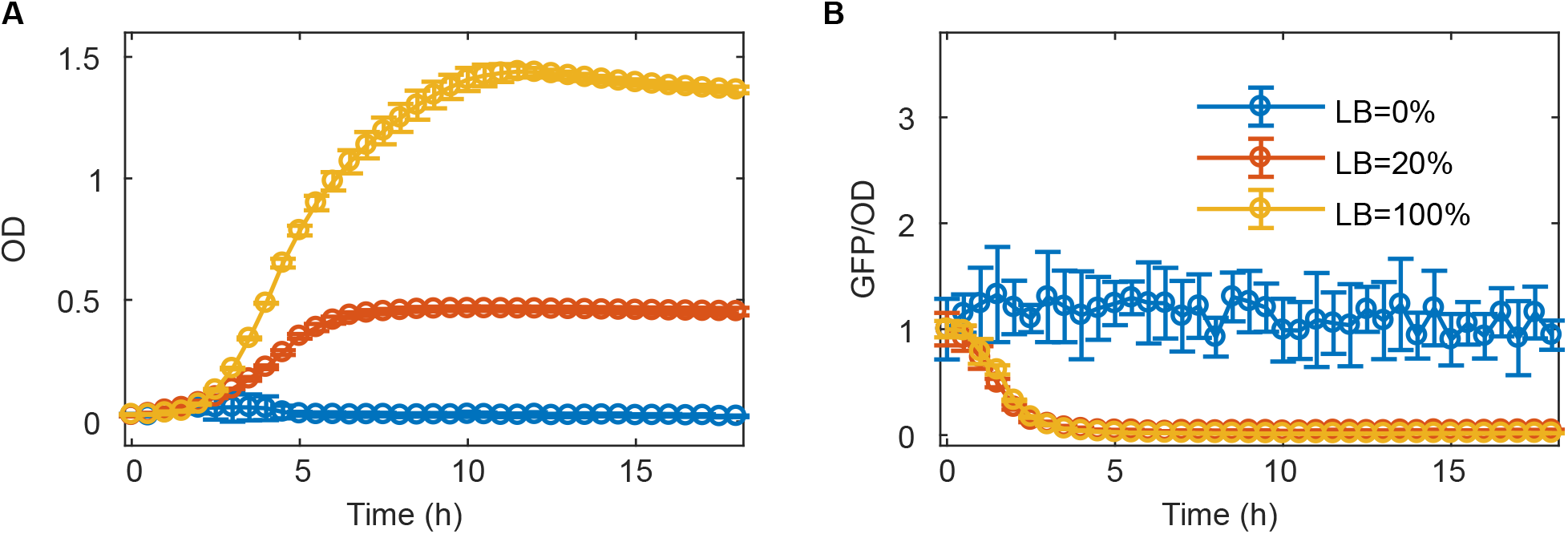
Memory loss of the SA circuit with moderate nutrient. (A-B) Dynamics of growth (Optical Density, OD) (A) and GFP/OD (B) after 1:100 dilution of activated cells into fresh medium without L-ara and three levels of LB fractions. The data with 100% LB and 0% LB are from previous work [18]. Data indicate mean±s.d. (n=4 biological replicates).

**Figure S2:**
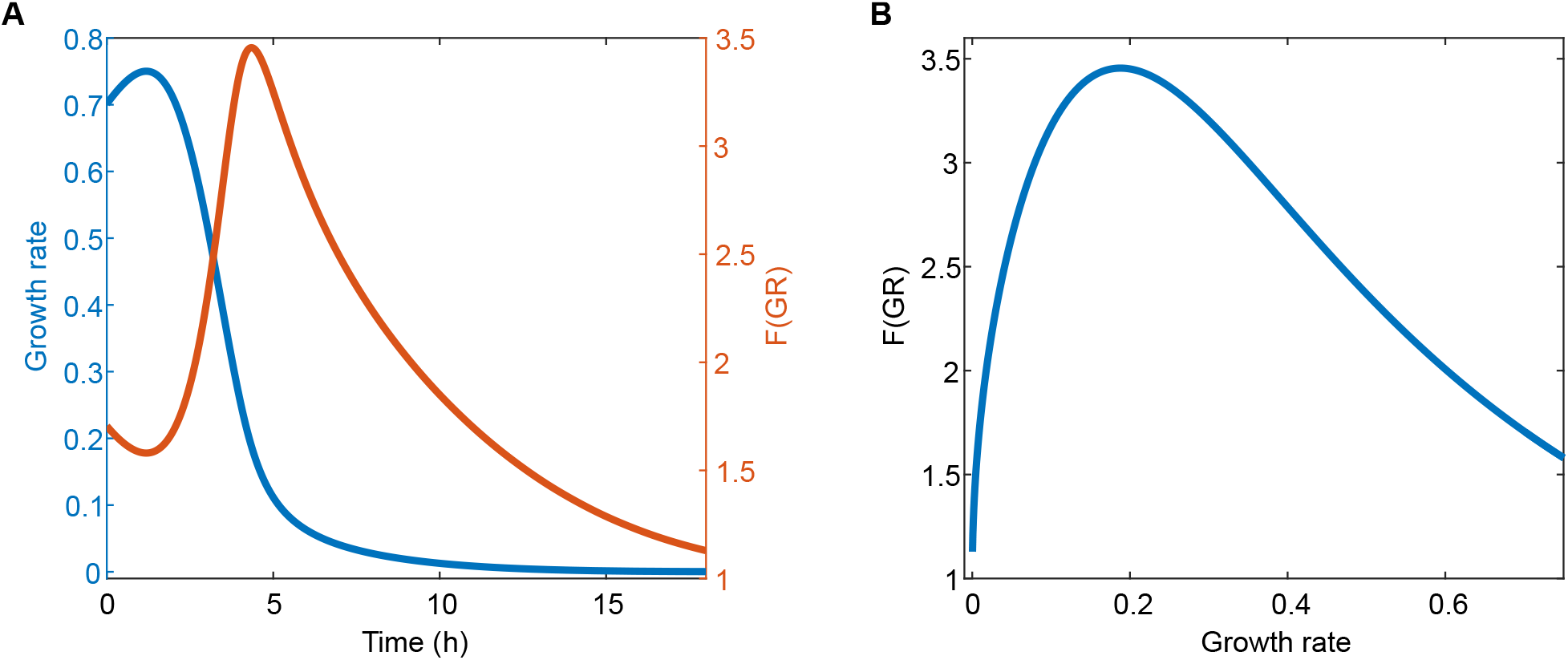
Fitting of the model to the dynamics of the host cell growth and circuit gene expression by considering a none (A), linear (B), monotonic (C) regulation of the production rete of the gene circuit by growth rate.

**Figure S3:**
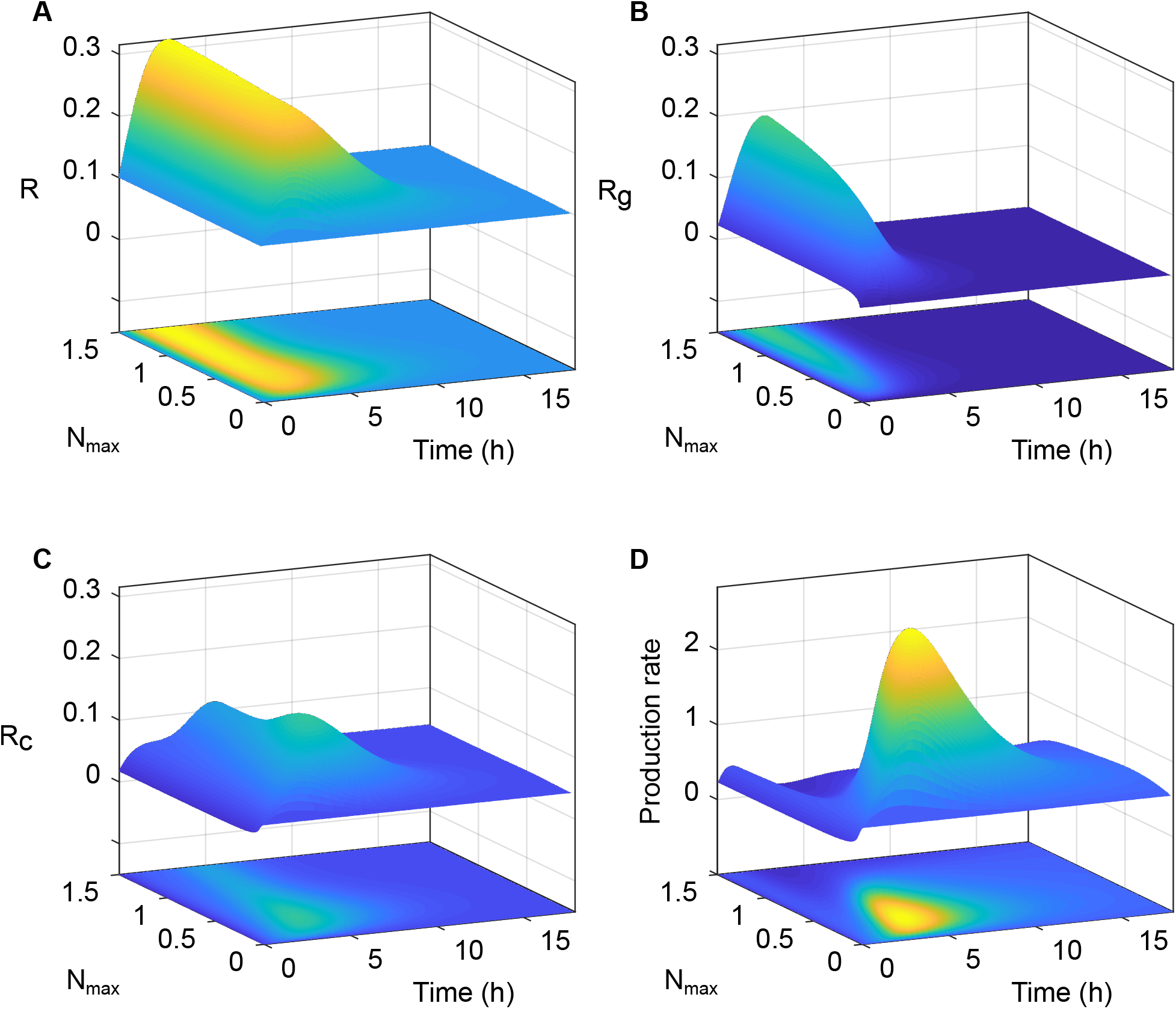
The dynamics of the total ribosome level (A), ribosome allocated to host cell growth (B), ribosome allocated to synthetic circuit (C), and the production rate of the circuit (D) with various carrying capacity (*N_max_*) in the SA circuit.

**Figure S4:**
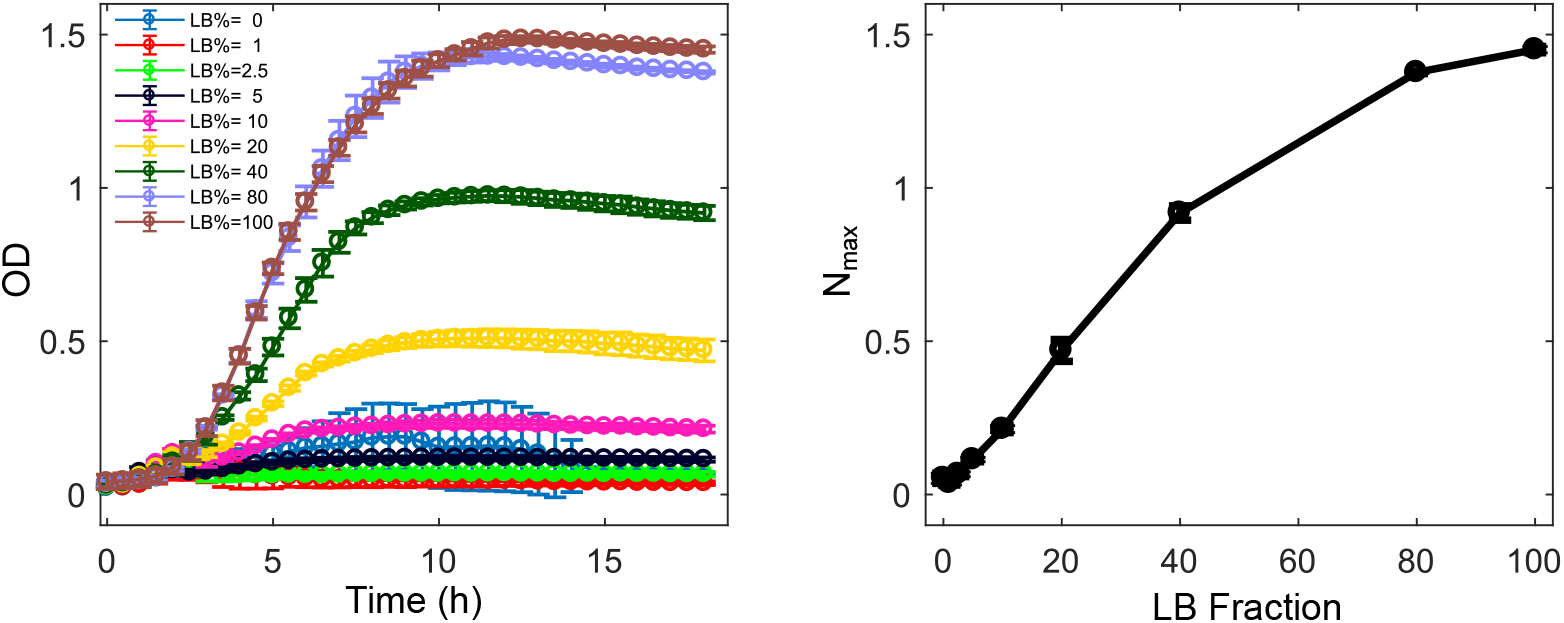
*N_max_* is a good parameter to represent the nutrient level in the theoretical analysis. (A) Dynamics of growth (OD) after 1:100 dilution of cells with activated self-activation switch into fresh medium with various LB fractions. The data for the LB=0, 20% and 100% used in Fig. 1 is repeated here for comparison. (B) The carrying capacity (*N_max_*) on the LB fraction in the culture medium. Data indicate mean±s.d. (n=4 biological replicates).

